# Deconvolution of Nascent Sequencing Data Using Transcriptional Regulatory Elements

**DOI:** 10.1101/2023.10.11.561942

**Authors:** Zachary Maas, Rutendo Sigauke, Robin Dowell

## Abstract

The problem of microdissection of heterogeneous tissue samples is of great interest for both fundamental biology and biomedical research. Until now, microdissection in the form of supervised deconvolution of mixed sequencing samples has been limited to assays measuring gene expression (RNA-seq) or chromatin accessibility (ATAC-seq). We present here the first attempt at solving the supervised deconvolution problem for run-on nascent sequencing data (GRO-seq and PRO-seq), a readout of active transcription. Then, we develop a novel filtering method suited to the mixed set of promoter and enhancer regions provided by nascent sequencing, and apply best-practice standards from the RNA-seq literature, using *in-silico* mixtures of cells. Using these methods, we find that enhancer RNAs are highly informative features for supervised deconvolution. In most cases, simple deconvolution methods perform better than more complex ones for solving the nascent deconvolution problem. Furthermore, undifferentiated cell types confound deconvolution of nascent sequencing data, likely as a consequence of transcriptional activity over the highly open chromatin regions of undifferentiated cell types. Our results suggest that while the problem of nascent deconvolution is generally tractable, stronger approaches integrating other sequencing protocols may be required to solve mixtures containing undifferentiated celltypes.

## 1. Introduction

One key problem of interest when studying transcription is the ability to capture the heterogeneity that exists in true biological samples.^1^ Bulk sequencing samples from cells are an aggregate across a cellular population, and thus average out differences between individual cells to capture only an ensemble profile of a given sample. Notably in the case of samples taken from tissues composed of heterogeneous constituent cells, any celltype specific differences are not necessarily discernible in the heterogeneous mixture of expression data.

To some extent, this problem has been at least partially solved in the context of RNA-seq with the emergence of single cell RNA-seq protocols which allow for RNA content at the level of the individual cell to be measured.^2^ However, the relatively high cost of sampling deeply limits the use of scRNA-seq in many contexts. Consequently, a great deal of work has been done to separate samples into constituent cell types *in silico*. This task is interchangeably referred to as deconvolution or microdissection. Deconvolution has been studied extensively in the context of both microarray data and in RNA-seq,^1,3–6^ but has seen only limited application to other high throughput genomic data.

Nascent transcription protocols^7,8^ are of particular interest for studies into transcriptional regulation.^9,10^ Nascent sequencing protocols profile active RNA Polymerase II activity, which captures enhancer associated RNAs (eRNAs), short unstable transcripts that are often associated with transcription factor binding sites.^11^ These eRNA transcript have proven to be highly informative markers of transcription factor activity.^9,10,12–16^ Unfortunately RNA-seq, whether bulk or single cell, does not capture enhancer associated transcripts due to the fact they are unstable and not polyadenylated.^11^ For this reason, the theoretical possibility of single cell measures of nascent transcription has tremendous potential for understanding regulation and transcription factor activity in key biological processes including development and disease progression.

Today, nascent sequencing protocols still operate only on the bulk level, largely because nascent protocols are relatively onerous, taking up to a week to process a set of samples.^7,8,17^ Because nascent protocols capture RNA production, many of the signals arise from lowly abundant, highly unstable RNAs.^11^ Furthermore, with current biochemical efficiencies, a single cell nascent sequencing protocol is likely infeasible, and thus deconvolution is needed to dissect nascent transcription profiles within tissues.

Nascent transcription data has relatively unique properties compared to RNA-seq. First, RNA-seq measures steady state mature, stable RNA levels which tend to be of relatively high abundance. In contrast, nascent sequencing protocols cover a much larger proportion of the genome (∼ 40% as opposed to ∼ 8%).^17^ The consequence is that the average sequencing depth per transcript is typically lower in nascent data, in spite of often sequencing samples to a higher depth. Second, many transcripts measured in nascent protocols are unannotated, lowly transcribed, unstable eRNAs (Figure 1A).^11,17^ In development, enhancer activities are the first changes detectable when a cell undergoes state change, suggesting their associated eRNAs have high potential as cell type markers.^18^ Furthermore, enhancer associated RNAs tend to be more cell type specific than protein coding genes.^19^ However, their low transcription levels lead to issues of reliable detection.^17^ Thus methods developed for RNA-seq must be appropriately adapted to use with nascent sequencing data.

**Figure 1.**
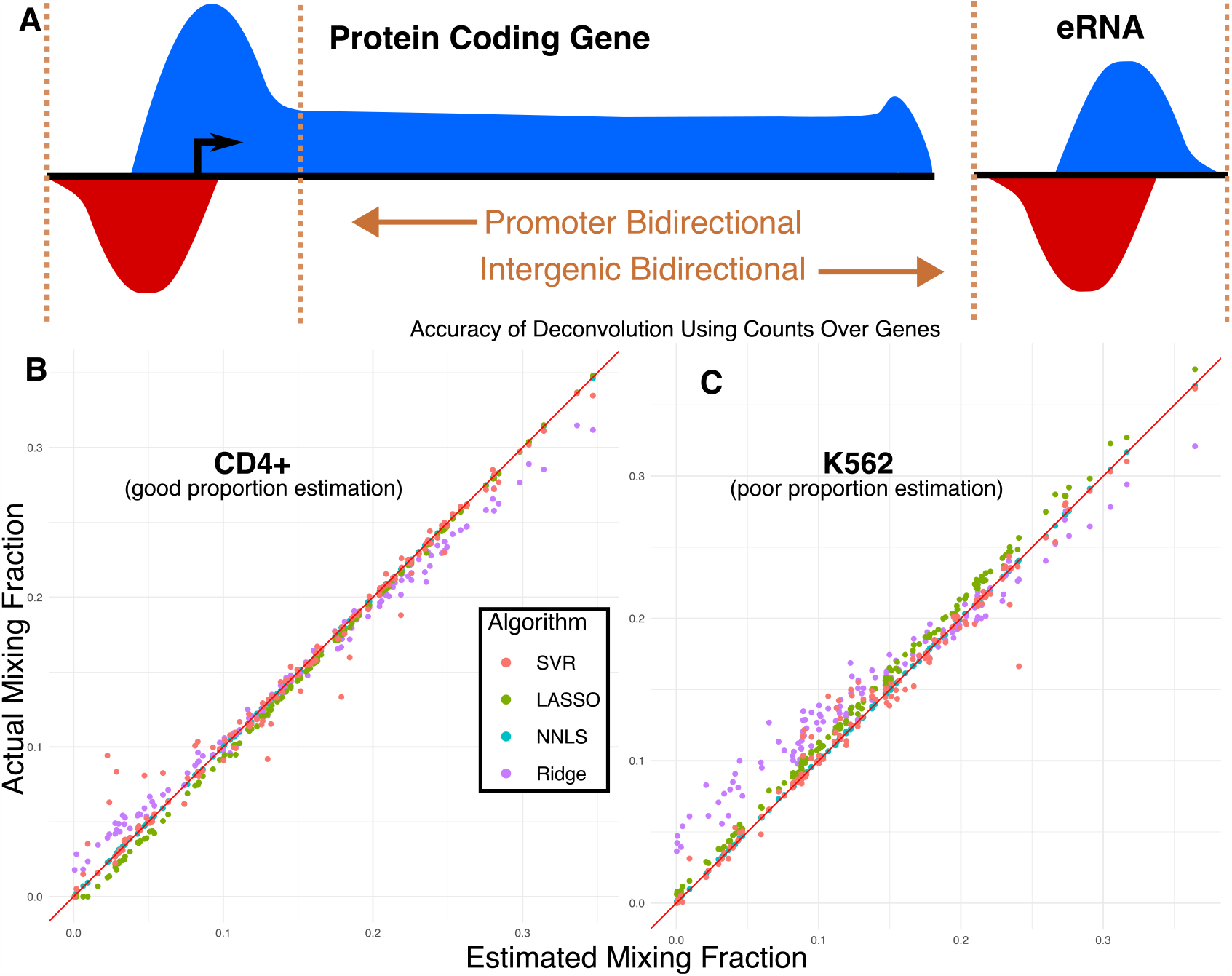
*A:* Nascent transcripts accumulate in a known bidirectional pattern around promoter sites as well as at enhancers.^7,36^ These bidirectional regions are counted by convention around 300bp from the site of RNA Polymerase initiation (roughly the center of the bidirectional).^9,10,36^ For annotated genes, we exclude the initiation peak by counting ±300 to the annotated transcription end site. *B:* Deconvolution was performed on random mixtures of cells from Table subsection 2.1. Some celltypes show highly accurate estimation of mixing proportion when doing deconvolution over all annotated genes, with most methods showing good linearity in their estimation. *C:* Other celltypes confound the regularized models used here, suggesting a systematic failure of regularization for proper estimation mixing proportion in this naive analysis. This failure appears to be more pronounced with L2 regularized methods and appears in all analyses conducted in this work, to some extent.

Here, we use standardized methods for supervised deconvolution to nascent sequencing data, applying a newly developed filtering technique to solve problems presented by nascent data in the deconvolution context. We show that deconvolution of nascent sequencing data works reliably, albeit with different model performance than in RNA-seq. We find that eRNAs present an informative set of information for deconvolution that can be inferred without a reference annotation. Furthermore, we find that undifferentiated celltypes confound deconvolution of nascent sequencing data, likely because their transcriptional expression resembles that of an aggregate of different differentiated celltypes.

## 2. Results

The problem of supervised deconvolution with sequencing data is formulated as follows: *Given sequencing samples from homogenous cell types and a heterogenous sample made up of those cell types, can we estimate the mixing proportions of those constituent cell types?* The problem of supervised (or partial) deconvolution is typically formulated as a linear system (Equation 1).^5,20^

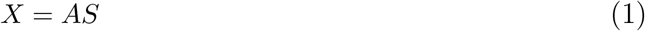

Here, X is a single-row matrix with one column per region of interest (ROI) (1 × *g*), A is a single row matrix with one column per reference homogenous cell type (1 ×*s*), and S is a matrix with one row per sample and one column per ROI (*s* × *g*). In most contexts, regions of interest (ROIs) correspond to annotated genes.

This is a overdetermined linear system, since the number of ROIs far exceeds the number of constituent cell types. Additionally, because these are biological values sampled from a noisy process, the key challenge is minimizing errors when solving the system. Most work in the literature has sought to solve the issues of this system in the context of RNA-seq or microarray^1,3–6,20–22^ data, with limited applications of this approach to other kinds of sequencing data.

For RNA-seq, a large variety of tools and approaches have been developed,^1,3–5,5,6,20–22^ which approach the problem using different models, constraints, and regularization approaches, as well as different ways to shrink the linear system. Many of these approaches claim to be the state-of-the-art, with most tools providing good performance. Consequently, we first examine the deconvolution problem on nascent sequencing using annotated genes and methods developed for RNA-seq.

### 2.1 Deconvolution on annotated genes

To evaluate existing deconvolution methods on nascent sequencing data, we first identified a number of high quality nascent sequencing data sets from a variety of cell types (see Table 2.1). Samples were processed using a standardized analysis pipeline^23^ which includes quality control, mapping and bidirectional transcript identification. These bidirectional transcripts originate from both gene start sites and regulatory elements such as enhancers (Figure 1A). The non-gene associated bidirectionals are often referred to as enhancer associated RNAs, or eRNAs.

**Table 2.1:** Samples used in this study.

| Study | GEO Accession | SRR | Cell Type |
| --- | --- | --- | --- |
| **Samples used in Figure 1–3** |  |  |  |
| Jiang 2018<sup>26</sup> | GSM3025555 | SRR6789175 | HCT116 |
| Fei 2018<sup>27</sup> | GSM3100195 | SRR7010982 | HeLA |
| Andrysik 2017<sup>28</sup> | GSM2296635 | SRR4090102 | MCF7 |
| Dukler 2017<sup>29</sup> | GSM2545324 | SRR5364303 | K562 |
| Zhao 2016<sup>30</sup> | GSM2212033 | SRR3713700 | Kasumi-1 |
| Danko 2018<sup>31</sup> | GSM3021718 | SRR6780907 | CD4+-T-cell |
| Chu 2018<sup>32</sup> | GSM3309955 | SRR7616132 | Jurkat-T-cell |
| **Samples added for Figure 4** |  |  |  |
| Core 2014<sup>33</sup> | GSM1480326 | SRR1552485 | GM12878 |
| Smith 2021<sup>34</sup> | GSM4214080 | SRR10669536 | ESC |
| Ikegami 2020<sup>35</sup> | GSM4207079 | SRR10601203 | BJ5ta |

As a first test, we examined only annotated protein coding genes to mimic deconvolution analysis typically done in RNA-seq. Notably, nascent data differs from RNA-seq in that splicing information is not present in nascent sequencing experiments, as RNA is collected pre-splicing. Furthermore, consistent with standards in nascent transcription analysis,^17^ we exclude the +300 initiation region of each gene when using featureCounts^24^ to count reads (see Figure 1A), as this avoids the 5′ bidirectional peak.

To simulate a mixed sample, we generated 128 randomly mixed samples by subsampling reads from each reference sample. Samples used for all *in-silico* experiments in this paper were mixed proportionally from raw reads using samtools,^25^ and are listed in Table 2.1. With these randomly mixed samples, we then performed supervised deconvolution using 4 different methods which are commonly discussed in the literature — Nonnegative-Least Squares Regression (NNLS), Ridge Regression, LASSO Regression, and *ϵ*-Support Vector Regression (SVR). For all methods tested, we apply a nonnegativity constraint (all mixing proportions must be at least zero) and a sum-to-one constraint (all mixing proportions must sum to one), as suggested in prior work.^1^ These constraints serve to make results from various deconvolution procedures interpretable as mixing weights for the linear deconvolution system. Code and supplemental materials for this project are available at https://github.com/Dowell-Lab/DeconvolutionNascent. We find that these methods provide generally good accuracy on deconvolution on our 128 randomly generated mixtures, although it appears that regularized methods perform more poorly than naive NNLS (Figure 1B,C) in certain celltypes across these mixtures. In this context, it appears that regularization does not improve accuracy at the cost of significant computational slowdowns relative to NNLS. Given these promising initial results, we next sought to shift the focus away from annotated genes to the unannotated bidirectional transcripts present at both promoters and enhancers.

### 2.2 Identifying bidirectionals as regions of interest

In addition to transcription at annotated genes, nascent transcription data contains bidirectional transcription at both promoters and regulatory elements. While annotated genes are widely studied and the typical target for this class of deconvolution algorithms, the study of enhancer associated RNAs is important for understanding the regulatory landscape of the cell. Various methods exist to identify sites of bidirectional transcription^36–39^ and to combine them across different samples.^10^ As such, bidirectionals are an additional region of interest that we now consider in our deconvolution framework.

To this end, we use a combined set of 485,688 bidirectionals, identified by Tfit and dReg within the Nascent-flow framework, capturing both enhancer RNAs and promoter regions, for all samples in Table 2.1.^36,38^ Notably, this system is significantly larger than the set of protein coding genes (approximately 490,000 vs 20,000). In this work, we use the following terminology in reference to subsets of this system — Bidirectionals refers to any site of RNA polymerase II initiation and generally includes both promoters and enhancers; any bidirectionals whose 5^*′*^ end (+/-300bp annotated TSS) overlaps an annotated 5′ gene in the RefSeq hg38 annotation is called a promoter; all other bidirectionals are called enhancers. Given the large size of this system, we next turn our attention to filtering the set of bidirectionals, to shrink the size of the overdetermined system to make deconvolution more computationally feasible.

### 2.3 Filtering methods are useful for shrinking the system

In traditional deconvolution contexts like microarray and RNA-seq, patterns of differential expression are often leveraged to shrink the system. For example, CIBERSORT^40^ uses an adaptive filtering method based on DESeq2 to find genes most indicative of specific celltypes. In the context of nascent sequencing data, however, tools like DEseq2 are problematic. The relatively low read coverage and cell type specificity of bidirectionals (e.g. inherent variability) leads DESeq2 to distrust these regions. To counter this, we developed a naive filtering scheme, selecting a fixed number of ROIs defined by the user for each homogenous reference sample where the reads for that sample were most different compared to all other samples. More formally, we define an algorithm for pruning the system of ROIs to a tractable level:

- Filter all ROIs to restrict them to regions where all celltypes have counts lower than the 99th percentile of reads in the sample. We do this to remove outliers whose extreme values could break the assumptions of a linear system.
- Generate transformed ratio *T* such that for each ROI (row), for each celltype (column), that entry is the log2 ratio of the count at that ROI over the maximum count for that ROI not in that celltype. This step generates a log2 transformed list of the ROIs that are the most specific to a single celltype.
- Order this list by the largest log2 ratio in any celltype in any ROI. Then, walk down this list keeping ROIs such that the number of ROIs for each celltype is approximately equal, up to some limit of elements. This generates a subset of the full system with the most celltype specific elements for each cell. The number of ROIs is approximate because the number of celltype specific elements varies per-celltype and can be exhausted at larger system sizes.

### 2.4 Most linear methods perform with high accuracy on synthetic nascent data

Given that bidirectional regions have distinct transcription characteristics compared to more robustly transcribed annotated genes, we first sought to assess deconvolution methods on the filtered bidirectional set. Using this set, we find that deconvolution achieves a high degree of accuracy (Figure 2A). Unexpectedly, we observe that across all sizes of system tested (including systems far in excess of the total number of genes in the human genome), non-negative least squares (NNLS) regression performs with the highest degree of accuracy. LASSO (L1 regularized linear regression) has a close second in performance. This is likely because LASSO regularization will only drop out cell types that are unlikely to be present in the mixture. In contrast, Ridge Regression (L2 regularized linear regression) performs worse than all other tested methods for most system sizes. Similarly, *ϵ*-Support Vector Regression (*ϵ*-SVR) with L2 regularization also performs relatively poorly compared to NNLS, but relatively well compared to Ridge regression. Despite these differences in accuracy, all models perform reasonably well on our synthetic mixtures, achieving accuracy to within a few percent on randomized mixtures. This is notable because these deconvolution methods perform well both on systems much smaller and much larger than those typically used for deconvolution of RNA-seq data.

**Figure 2.**
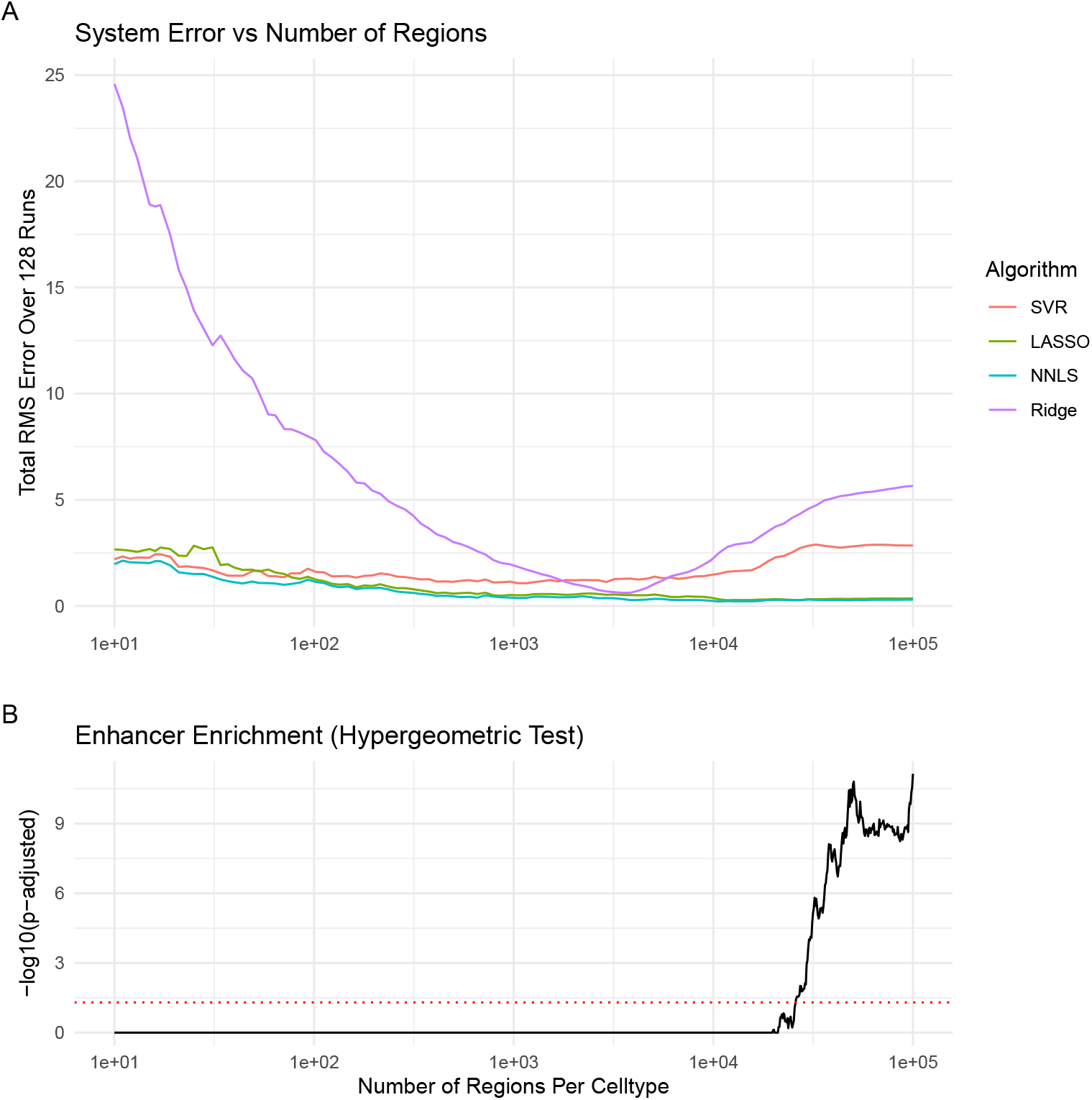
*A:* Models were tested using standard library implementations on a set of 128 randomly generated sets of mixing parameters. Each model was tested on 100 different subsets of ROIs selecting 10^*n*^-many points for *n* ∈ [1, 5] using linear spacing between subsequent *n*. Most models perform well in the intermediate region of 10^3^–10^4^ points selected per-sample, but diverge outside of that regime. For each set of ROIs selected, the same 128 randomly generated sets of mixing parameters were used as in Figure 2. We observe that for essentially all points, NNLS outperforms more complex models. *B:* To understand the selection process of our subsetting algorithm, we tested whether enhancers were selected from the full ROI set at a greater rate than would be expected by random. To do so, we performed a hypergeometric test with Bonferroni correction over all trials of our ROI subsets. We observe that for smaller system sizes the enhancer/promoter sampling ratio does not differ dramatically from that expected by random sampling. When the system size increases, enhancers become preferentially selected over promoters (*p <* 0.05), but this increase in the rate of enhancer selection does not correlate with the accuracy of any model.

Interestingly, we find our subsetting method consistently selects a mixture of enhancers and promoters that does not significantly differ from the distribution expected by random chance (Figure 2B). Consequently, this procedure captures mostly eRNAs and not promoters, since the number of eRNAs far outnumbers the number of promoters. This suggests that certain enhancer-driven regulatory elements are highly informative in identifying celltype.

We next sought to determine which ROIs were most informative to the deconvolution problem. To answer this question, we utilized NNLS, the best performing method in our prior tests. Using NNLS, we compared the performance on bidirectionals (as in Figure 2A), to annotated genes (as in Figure 1B,C) and a combination of these features – selected using our region filtering approach (Figure 3). We find that these methods achieve high accuracy for both genes and bidirectionals across a number of system sizes, with somewhat reduced accuracy when combining these two sets of ROIs. This reduction in accuracy could be a result of colinearity in the combined set of ROIs, as some bidirectionals may be intronic and thus they are not a strictly non-overlapping set relative to annotated genes.

**Figure 3.**
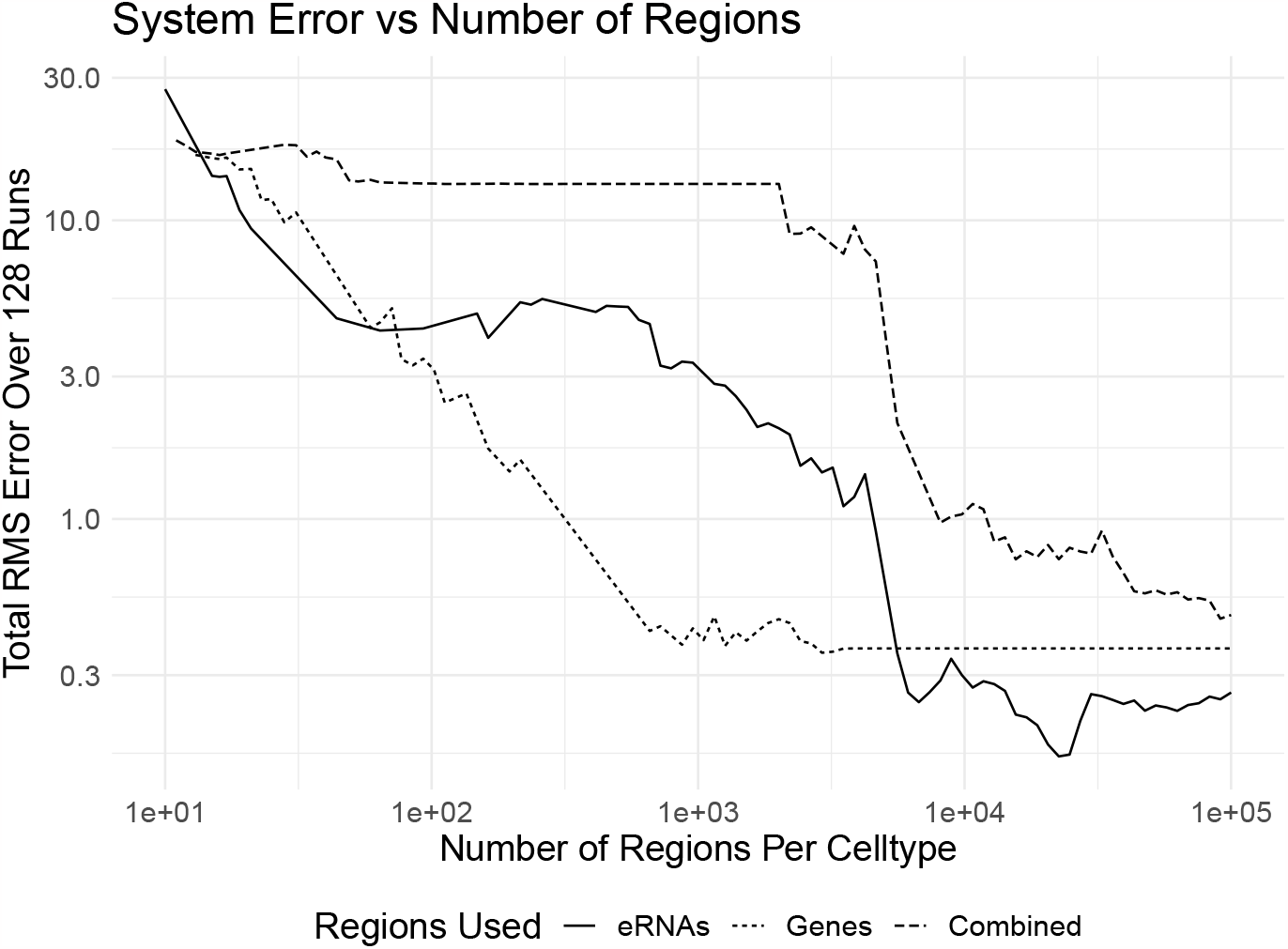
To compare the maximum theoretical accuracy of our system, we conducted the same analysis as in Figure 2 using either the region sets of bidirectionals, annotated genes, or a combination of the two, performing the same subsetting procedure as before. We observe that at smaller region sizes using genes alone provides a higher degree of accuracy than just bidirectionals, but that at larger sets of ROIs bidirectionals alone can achieve a higher absolute degree of accuracy. Somewhat unexpectedly, the combination of both sets of regions performs more poorly than each separate subset. Note that as system size increases, the accuracy of the set using annotated genes reaches a constant level purely because the total size of that system is exhausted by virtue of being an order of magnitude smaller than that of the bidirectionals or combined set.

For the data tested and the size of system used, we found that certain methods in the literature were prohibitively slow for the large linear systems we tested. For example, a *ν*support vector regression (*ν*-SVR) approach as suggested by CIBERSORT^40^ was too computationally expensive to test or benchmark reliably, taking more than 24 hours to do deconvolution on a single mixture of cells at large system sizes (approximately 100k ROIs or more). Due to these poor scaling characteristics, we instead chose to use an optimized implementation of the primal version of *ϵ*-SVR. This was chosen instead of a dual formulation to maintain computational tractability for the large number of samples relative to the number of features. In the context of nascent sequencing data, NNLS is likely the best model to use based on our benchmarking.

### 2.5 Undifferentiated celltypes confound deconvolution of mixtures

In the course of testing our model, we observed that certain celltypes strongly confounded all deconvolution models tested when using bidirectionals. To understand this puzzling behavior, we examined deconvolution in the presence and absence of these cell types. To do deconvolution of this system, we generated a titration curve, mixing celltypes from distinct separate mixing proportions into equivalent proportions for all celltypes.

We observed that both ESC cell lines and BJ5TA cell lines caused deconvolution to fail (Figure 4A,B). Specifically, inclusion of either cell line results in an overestimation of the mixing proportion for those cell types. We carefully examined these two cell lines to identify distinguishing features relative to the other cell lines.

**Figure 4.**
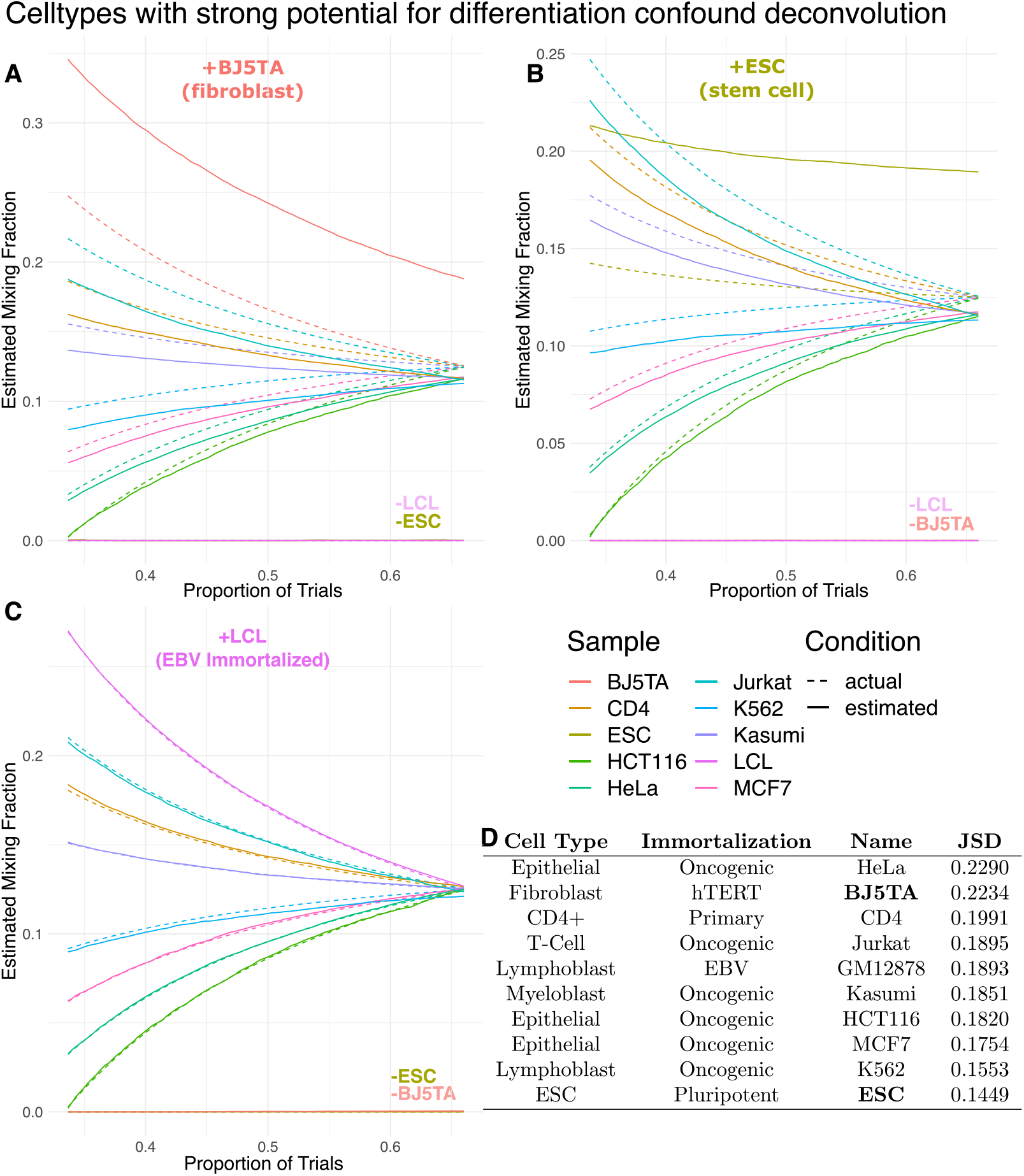
To interrogate the effect of undifferentiated and partially differentiated celltypes on the performance of deconvolution, we performed a titration experiment, estimating mixing parameters for 100 different mixtures of celltypes as mixing proportions were taken from maximally separated to equivalent. For each trial *n*, the mixing proportions are equally spaced points in 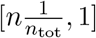 that are then rescaled to sum to one. Each subset (A,B,C) was generated by holding out one celltype from the full mixture and renormalizing the adjusted mixing proportions to sum to one. *A,B:* Adding either BJ5TA or ESC cells into the mixture causes a higher-than-true proportion of those cells to be estimated. Neither of these cell lines are terminally differentiated. *C:* Addition of EBV immortalized LCL cells into the mixture does not result in failure of the deconvolution model, suggesting that the observed failures are not a function of how cells were immortalized. *D:* To understand if this failure could be attributed to celltype specificity, we calculated the mean Jensen-Shannon Divergence for each sample compared to all others. The pluripotent ESC cells show the lowest celltype specificity while the partially differentiated BJ5TA cells show the highest celltype specificity, with the exception of HeLa cells.

To determine whether the number of cell lines or cell line immortalization differences could be the source of the problem, we added lymphoblastoid cell lines immortalized (LCL) by EBV. Notably, LCLs do not confound the model and show excellent performance (Figure 4C). Both ESC (embryonic stem cells) and BJ5TA (fibroblast derived) are non-terminally differentiated and non-oncogenic (Figure 4D). Furthermore, we see that even without regularization, NNLS successfully removes non-present celltypes (Figure 4A-C), meaning that undifferentiated cell-types will not be inferred in the mixing proportion if they are not present at all in the mixture. Furthermore, regularization techniques are not required to accomplish this removal of celltypes that are absent.

One alternative hypothesis to the source of this problem is that heterogeneity in the population of undifferentiated celltypes is the source. However, this would suggest that more heterogeneous cell populations should perform worse in deconvolution, as should cells from similar tissue types. Yet based on this data, this seems unlikely, given that both CD4+ and Jurkat cells, both peripheral blood mononuclear cells (PBMC) derived, are present in the mixture and are successfully estimated by our models. Since the addition of a lymphoblast cell line immortalized using EBV (GM12878) does not result in system failure in the same way that is observed with the non-differentiated cell-lines, we suspect that differentiation is the key issue here as opposed to heterogeneity. Our work suggests that undifferentiated or partially differentiated cell types pose a key challenge to the deconvolution of nascent sequencing data when using enhancers because their regulatory profile, particularly that of their enhancer regions, resemble an ensemble profile of multiple differentiated celltypes. In support of this, the problem does not seem to occur when using genes alone, suggesting that undifferentiated cells may lack the same level of specificity at bidirectionals as terminally differentiated cell types.

Our results suggest either very low or very high celltype specificity when looking at these samples’ bidirectional ROIs (Figure 4D). When looking at the mean Jensen Shannon Divergence for each sample compared to all others, we observe that our undifferentiated cell lines are either the least specific (ESC) or the most specific (BJ5TA). Although HeLa cells show the highest degree of celltype specificity by this measure, HeLa cells are not representative of human cells, exhibiting notably different expression patterns^41^ which would lead to a high degree of cell type specificity. Past work has shown that ESC cell lines have genome-wide transcriptional hyperactivity^42^ that narrows as differentiation progresses. Additionally, work in hematopoetic cells has suggested that these undifferentiated cell lines are characterized by a high degree of fluidity in chromatin modification.^43^ More work is required to definitively establish that differentiation is the source of the breakdown of deconvolution in this system, and will likely require significant work outside the scope of this preliminary study.

## 3. Conclusion

This work is the first to examine supervised deconvolution of heterogenous mixtures of nascent sequencing data. Deconvolution is an essential tool for the study of heterogenous samples, whether cell lines or tissues. While most work on deconvolution of heterogenous samples has moved on to focusing on single cell protocols, a single cell nascent sequencing protocol currently seems infeasible. Thus, nascent sequencing is limited to bulk experiments, which appear to be reliably separable by supervised deconvolution. We present here the use of nascent sequencing data as a testbed for this supervised deconvolution problem. We integrate best practices from the literature and develop new techniques to handle characteristics in nascent sequencing data where assumptions from the RNA-seq deconvolution literature do not hold.

To benchmark various deconvolution algorithms, we first developed a new algorithm to filter ROIs to only use regions with the most celltype specific expression. We find that this selection process does not preferentially select enhancer or promoter ROIs. That said, the number of enhancer associated bidirectionals far exceeds annotated genes, providing ample features from which to select regions of interest. Our proposed algorithm is simple, fast, and reliable, and establishes a strong first basis for the development of more specific ROI filtering tools for nascent deconvolution.

Using this algorithm, we compared standard methods used for solving the deconvolution problem. Specifically, we tested NNLS, Ridge, LASSO, as well as *ϵ*-SVR. We found that all methods reliably separate the nascent deconvolution system, with L2-regularized methods achieving comparatively poor performance to NNLS. Furthermore, we found that even a simple method like NNLS could reliably eliminate celltypes that were not present in the sample, suggesting regularization is not necessary for solving the deconvolution problem here. While we find that both annotated genes and bidirectionals can achieve high accuracy in supervised deconvolution (with bidirectionals having an edge in absolute accuracy), it is worth emphasizing that bidirectionals are distinctly advantageous in that they are annotation-independent and discovered *de-novo* for each sample.

We show that the addition of undifferentiated samples to a nascent deconvolution system results in highly skewed mixing estimates, with undifferentiated celltypes predicted as far more likely than their actual frequency in the mixture. One possible reason for this is that undifferentated celltypes tend to show regulatory patterns akin to a combination of the regulatory patterns of each constituent celltype. It appears to be a necessary condition for some amount of the undifferentiated celltype to be present in the mixture in order for the system to fail.

One key issue in this work is the lack of availability of diverse high quality nascent sequencing data to perform simulations against. Although a large amount of nascent sequencing data is available and published, the number of cell types available is somewhat limited. Protocols aimed at extending run-on sequencing to a broader base of samples, such as ChRO-seq^44^ show promise in alleviating this bottleneck. Importantly, many of the earliest nascent data sets lacked replicates – which excluded their usage here. Data quality and availability is often a limiting factor in computational studies, and this work is not an exception to that rule.

In this work, and generally for the supervised deconvolution problem, we assume that all cells in a sample are taken from an approximately homogeneous population. This is sometimes a reasonable assumption but is often not. One future frontier that could be highly beneficial to this project is the incorporation of single cell ATAC-seq (scATAC) as a secondary source of information to augment bulk nascent sequencing data. scATAC combines the chromatin accessibility readout provided by ATAC-seq (indicative of regions open to transcription) with the cell-specific information provided by modern single cell sequencing protocols. Tools are already well defined for clustering single cell sequencing data into constituent cell types, as individual cells can typically be separated using dimensionality reduction methods like PCA, tSNE, or UMAP.^45,46^ Because transcription occurs in regions of open chromatin, which is what ATAC-seq measures, mixing fractions and celltype specific transcripts could be estimated more reliably using combined data from both protocols. Future work combining pairing single cell ATAC-seq data and nascent sequencing data could leverage techniques used by existing tools^21^ to do deconvolution on a more granular level for individual samples, providing a strong complementary tool to the bulk deconvolution discussed here. While single cell approaches remain comparatively expensive, this combination would be a powerful tool for looking at transcriptional regulatory networks at the level of sub populations of samples.

Nascent sequencing is a powerful tool for the assessment of transcriptional regulatory networks, and when paired with deconvolution tools will also facilitate deeper understanding of those regulatory networks in heterogeneous cell populations. Leveraging a transcription oriented sequencing approach instead of an expression oriented (e.g. steady state) one provides myriad benefits — more thorough coverage of the genome, understanding of regulatory elements, and a deep view of underlying transcriptional dynamics — all of which can be integrated with different sequencing protocols to great effect. Supervised deconvolution represents an important preliminary foothold into this space, and this work shows that nascent sequencing data is well suited for that class of problems.

